# TGFBR2^High^ mesenchymal glioma stem cells phenocopy regulatory T cells to suppress CD4+ and CD8+ T cell function

**DOI:** 10.1101/2025.01.07.631757

**Authors:** Amanda L Johnson, Harmon S. Khela, Jack Korleski, Sophie Sall, Yunqing Li, Weiqiang Zhou, Karen Smith-Connor, Hernando Lopez-Bertoni, John Laterra

**Author notes:** **Corresponding Authors:** *Hernando Lopez-Bertoni, Ph.D.*, Johns Hopkins University School of Medicine and Hugo W. Moser Research Institute at Kennedy Krieger, 707 North Broadway, Baltimore, MD 21205, USA., *John Laterra, MD, PhD.,* Johns Hopkins University School of Medicine and Hugo W. Moser Research Institute at Kennedy Krieger, 707 North Broadway, Baltimore, MD 21205, USA.

## Abstract

Attempts to activate an anti-tumor immune response in glioblastoma (GBM) have been met with many challenges due to its inherently immunosuppressive tumor microenvironment. The degree and mechanisms by which molecularly and phenotypically diverse tumor-propagating glioma stem cells (GSCs) contribute to this state are poorly defined. In this study, our multifaceted approach combining bioinformatics analyses of clinical and experimental datasets, single-cell sequencing, and molecular and pharmacologic manipulation of patient-derived cells identified GSCs expressing immunosuppressive effectors mimicking regulatory T cells (Tregs). We show that this **I**mmunosuppressive **T**reg-**L**ike (ITL) GSC state is specific to the mesenchymal GSC subset and is associated with and driven specifically by TGF-β type II receptor (TGFBR2) in contrast to TGFBR1. Transgenic TGFBR2 expression in patient-derived GBM neurospheres promoted a mesenchymal transition and induced a 6-gene ITL signature consisting of CD274 (PD-L1), NT5E (CD73), ENTPD1 (CD39), LGALS1 (galectin-1), PDCD1LG2 (PD-L2), and TGFB1. This TGFBR2-driven ITL signature was identified in clinical GBM specimens, patient-derived GSCs and systemic mesenchymal malignancies. TGFBR2^High^ GSCs inhibited CD4+ and CD8+ T cell viability and their capacity to kill GBM cells, effects reversed by pharmacologic and shRNA-based TGFBR2 inhibition. Collectively, our data identify an immunosuppressive GSC state that is TGFBR2-dependent and susceptible to TGFBR2-targeted therapeutics.

## Introduction

The GBM tumor microenvironment (TME) is defined by low infiltration of anti-tumor immune cells, high prevalence of T cell exhaustion, and relatively high numbers of suppressive, pro-tumor immune cell infiltrates (1, 2). Tumor cell-intrinsic mechanisms leading to immunosuppressive TMEs are increasingly recognized as barriers to anti-tumor immunity and immunotherapy. These escape mechanisms are utilized by glioma stem-like cells (GSCs) to avoid recognition by the immune system and allow for continued tumor growth (3–5). GSCs can regulate the immune TME by producing factors that recruit immunosuppressive cells and inhibit cytotoxic T cells (5).

Subsets of GSCs possess unique phenotypic and immune-modulatory traits (6–11). Emerging evidence indicates that tumor cell phenotypic transitions dynamically contribute to establishing and maintaining the immunosuppressive TME in GBM (12, 13) and recent observations suggest that GBM dynamically adapts to different modes of immunotherapy by remodeling the tumor cell subtype composition (14), with mesenchymal transitions being of particular importance (12, 15). Which cell fate-determining events contribute to immune evasion in GBM and what aspects of these tumor cell transitions are amenable to therapeutic intervention remain unknown.

One critical mediator of immune regulation in the TME is TGFβ, a cytokine secreted by various cell types with context-dependent immune-regulatory functions (16). In GBM, TGFβ suppresses anti-tumor immune cells (e.g., T cells, dendritic cells) and promotes pro-tumor immune cells (e.g., tumor-associated macrophages, microglia, and regulatory T cells) (17). TGFβ signaling is also a well-established driver of stem-like traits and epithelial-to-mesenchymal transition (EMT) in GBM and other malignancies (17, 18). Canonical TGFβ signaling is initiated by ligand binding to the type II receptor (TGFBR2) that phosphorylates the type I receptor (TGFBR1) prior to downstream activation of the transcriptional regulators Smad2 and Smad3 by phosphorylation. TGFβ receptors are serine-threonine kinases with a variety of potential interacting proteins and downstream signaling effectors (19) that can signal independent of each other (20). The distinct roles of TGFBR1 and TGFBR2 in cancer, especially as they pertain to immune regulation in GBM, remain undetermined.

The goal of this study is to further understand how stem-cell driving events contribute to the cell-intrinsic immunosuppressive phenotype of GBM cells. By combining molecular manipulation of GBM cells, single-cell sequencing, computational analyses, and tumor-immune cell co-culture systems, we identified a novel TGFBR2-driven phenotype in GSCs with molecular parallels to regulatory T cell (Treg) fate induction. We show that mesenchymal-like GBM neurospheres enriched for GSCs (mGSCs) express high levels of TGFBR2 and are capable of repressing CD4+ and CD8+ T cell viability and function *in vitro*. Moreover, TGFBR2 inhibition reversed this immunosuppressive tumor cell phenotype by reducing CD8+ T cell exhaustion and enhancing CD4+ and CD8+ T cell-mediated tumor cell killing. These findings identify a novel mechanism of GSC immunosuppression and highlight the potential applicability of anti-TGFBR2-specific therapeutics for augmenting immunotherapy in GBM.

## Results

### Oct4 and Sox2 induce a TGFBR2-related mesenchymal shift in GBM cells

To explore clinically relevant Oct4/Sox2-induced reprogramming events in GBM, we began by conducting bulk RNA sequencing (RNA-seq) on patient-derived neurospheres with and without transgenic co-expression of Oct4 and Sox2, reprogramming transcription factors shown by us and others to induce tumor-propagating GSC phenotypes in GBM cells (5, 21–24). We then cross-referenced the resulting differential gene expression data to transcripts upregulated in clinical GBM specimens compared to non-tumor tissue. This analysis identified a collection of genes both induced by Oct4 and Sox2 and enriched in GBM compared to non-tumor brain tissue (**Fig. 1A**). Furthermore, 25 of these genes were found to be significantly upregulated in mesenchymal GBMs versus other molecular subtypes and of these genes TGFBR2 ranked highest (**Fig. 1A**; genes in red). Single-cell RNA sequencing (scRNA-seq) analysis conducted on patient-derived GBM neurospheres enriched for GSCs -/+ transgenic Oct4/Sox2 shows a shift towards the MES-like cell state and away from OPC-like, NPC-like and AC-like neurodevelopmental phenotypes (**Fig. 1B**) (25). Consistent with these transcriptomic signatures, Western blot analysis showed that Oct4/Sox2 expression increased mesenchymal protein markers Slug, Vimentin, and CD44 concurrent with decreased expression of the proneural marker CD133 (6, 7) (**Fig. 1C**). TGFBR2 was also most strongly associated with MES-like GBM cells relative to other neurodevelopmental subtypes (**Fig. 1D**), consistent with its upregulation by Oct4/Sox2 and association with clinical mesenchymal GBM (**Fig. 1A**). Western blot analysis confirmed upregulation of both TGFBR2 and phosphorylated TGFBR1, a surrogate for activated TGFβ signaling, in neurospheres expressing transgenic Oct4 and Sox2 (**Fig. 1E**). Moreover, Smad2/3 transcriptional targets were induced by Oct4/Sox2 preferentially in mesenchymal-like neurospheres (**Fig. 1F**), and qRT-PCR analysis confirmed upregulation of a subset of Smad2/3 target genes by Oct4/Sox2 co-expression (**Fig. 1G**). Further analyses of clinical, patient-derived xenograft, and primary GSC transcriptomic data revealed a consistent and strong positive correlation between TGFBR2 expression and the mesenchymal marker CD44 across these diverse datasets, contrasting the weak and negative correlations with the proneural marker CD133 (*PROM1*) (**Fig. 2A**). Notably, TGFBR1 expression did not positively correlate with CD44 across three complementary clinical, PDX and primary GSC datasets (**Fig. 2B**). Separating these same datasets into CD44 high/low showed that Smad2/3 transcriptional targets are enriched in CD44-high samples (**Fig. 2C**). These correlations were confirmed by Western blot analysis showing higher endogenous levels of TGFBR2 in mesenchymal patient-derived neurospheres compared to more classical neurospheres (**Fig. 2D**). Together, these observations predict that TGFBR2 signaling plays a prominent role in the mesenchymal transition driven by reprogramming events initiated by Oct4 and Sox2 in GBM.

**Figure 1.**
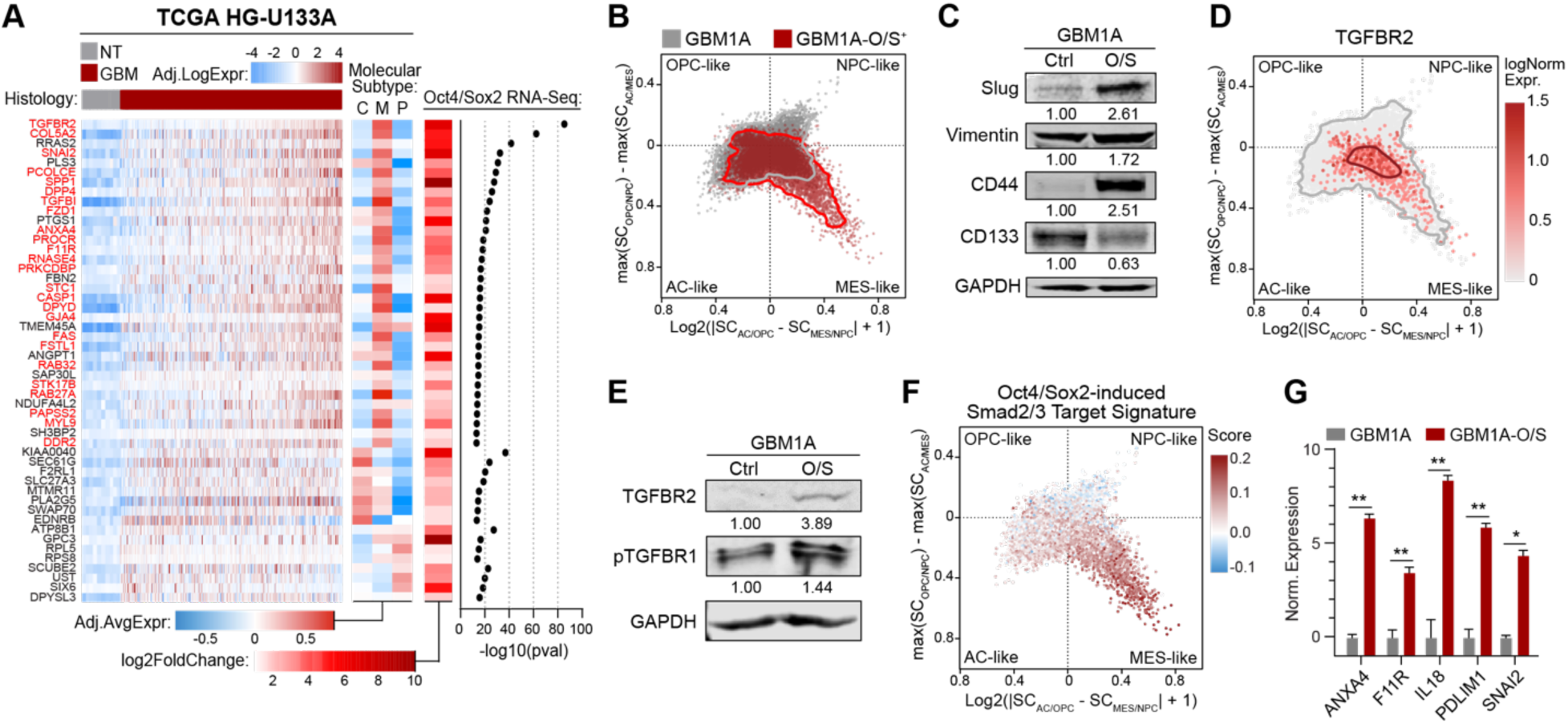
Oct4 and Sox2 drive a mesenchymal-like shift in GSCs associated with activation of TGFBR2 signaling. **(A)** Heatmap (left) showing expression of genes upregulated in IDH-wt GBM versus non-tumor (NT) brain tissue, enriched in mesenchymal GBMs (middle), and induced by transgenic co-expression of Oct4/Sox2 in patient-derived GBM neurospheres (right). Genes in red are significantly enriched in mesenchymal GBMs compared to classical and proneural. **(B)** Cell state plot of GBM1A neurospheres -/+ transgenic co-expression of Oct4/Sox2 (O/S). **(C)** Western blot analysis showing expression of mesenchymal driver Slug, mesenchymal markers Vimentin and CD44, and proneural marker CD133. Numerical values represent band signal intensity relative to parental (Ctrl) cells and normalized to GAPDH. **(D)** Cell state plot showing TGFBR2 expression in GBM1A and GBM1A-O/S neurospheres. **(E)** Western blot showing expression of TGFBR2 and phospho-TGFBR1 in GBM1A cells -/+ co-expression of Oct4/Sox2. Numerical values represent band signal intensity relative to parental (Ctrl) cells and normalized to GAPDH. **(F)** Cell state plot showing expression Smad2 and/or Smad3 transcriptional targets (GSE11710) induced by Oct4/Sox2 in GBM1A cells -/+ O/S. **(G)** qRT-PCR analysis comparing expression of a subset of Smad2/Smad3 targets in GBM1A cells -/+ O/S. Data are shown as mean ± SD. Statistical significance was calculated using Student’s T-test in panel G. *p<0.05, **p<0.01.

**Figure 2.**
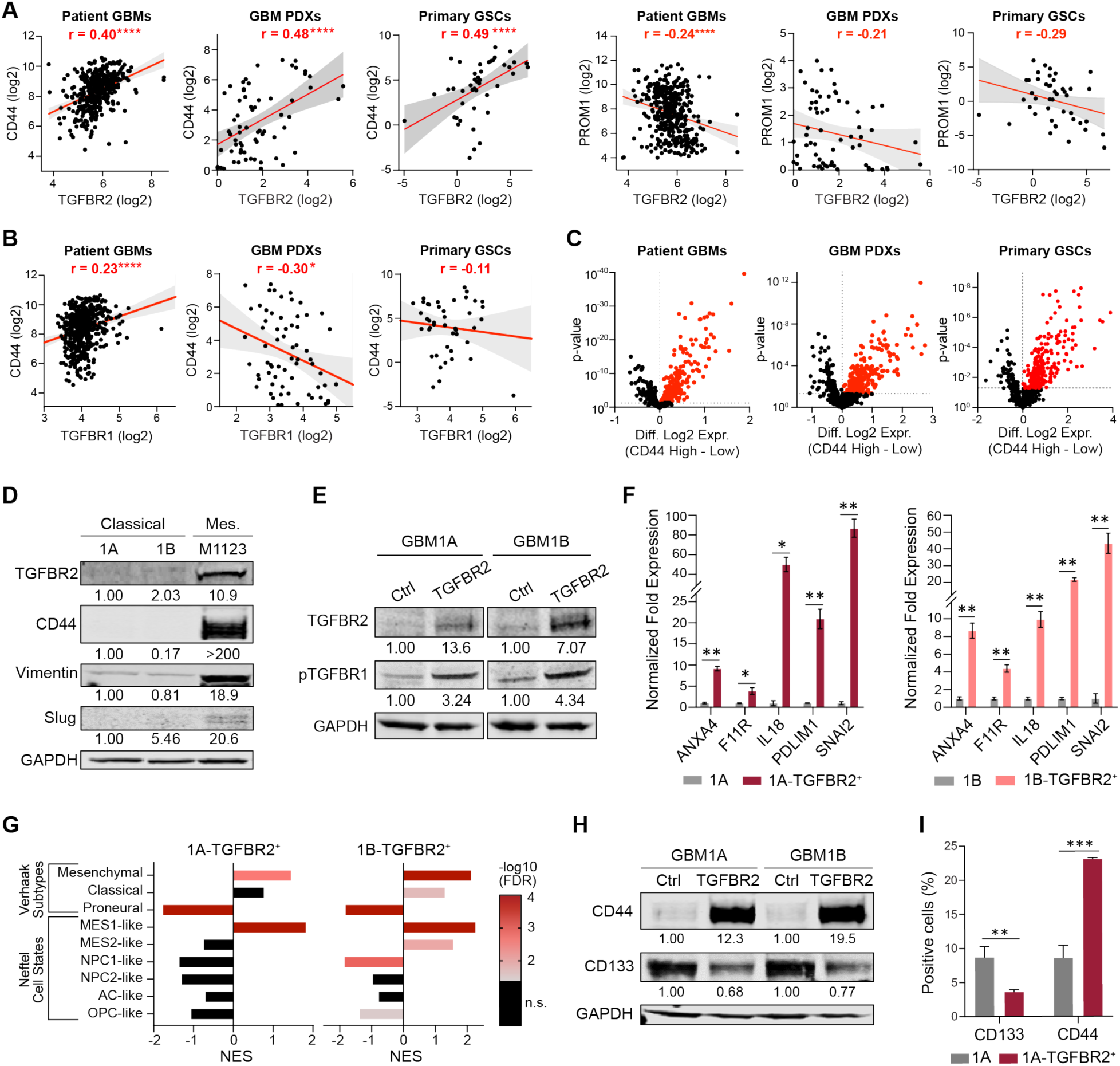
TGFBR2 is sufficient to induce a mesenchymal shift in GSCs. **(A)** Pearson correlation plots comparing mRNA expression of CD44 (left) and PROM1 (right) to TGFBR2 in GBM patient specimens (TCGA HG-U133A), GBM patient-derived xenograft (PDX) lines and primary patient-derived GSCs. **(B)** Pearson correlation plots comparing mRNA expression of CD44 to TGFBR1 in GBM patient specimens, GBM PDXs, and primary GSCs. **(C**) Volcano plots showing expression of Smad2/3 transcriptional targets in CD44 high vs. CD44 low samples from GBM patient specimens, GBM PDXs, and primary GSCs. **(D)** Western blot comparing expression of TGFBR2 and mesenchymal markers CD44, Vimentin, and Slug in GBM1A (1A), GBM1B (1B), and M1123 patient-derived neurospheres. Numerical values represent the band signal intensity relative to GBM1A and normalized to GAPDH. **(E)** Western blot showing expression levels of FLAG (TGFBR2) and phospho-TGFBR1 in GBM neurospheres -/+ transgenic FLAG-tagged TGFBR2. Numerical values represent the band signal intensity relative to the parental line (GBM1A or GBM1B) and normalized to GAPDH. **(G)** qRT-PCR analysis showing expression of a subset of Smad2/3 transcriptional targets in GBM neurospheres -/+ transgenic TGFBR2. **(G)** GSEA of GBM molecular subtypes from TGFBR2-induced transcriptomes. Color of bars represent the false discovery rate (n.s. = not significant). **(H)** Western blot showing CD44 and CD133 protein levels in neurospheres -/+ transgenic TGFBR2. Numerical values represent the band signal intensity relative to parental (Ctrl) cells and normalized to GAPDH. **(I)** Flow cytometry analysis to measure CD44+ and CD133+ cell populations following transgenic expression of TGFBR2 in GSCs. Data in panels F and I are shown as mean ± SD. Statistical significance was calculated using Pearson’s correlation in panels A and B and Student’s T-test in panels C, F, and I. *p<0.05, **p<0.01, ***p<0.001.

To test the hypothesis that TGFBR2 signaling is sufficient to drive a mesenchymal transition, we expressed transgenic TGFBR2 in classical GBM1A and GBM1B neurospheres that endogenously express low levels of TGFBR2 (**Fig. 2D**). Transgenic expression of TGFBR2 was sufficient to activate TGFβ signaling, as measured by TGFBR1 phosphorylation (**Fig. 2E**) and expression of down-stream Smad2/3 transcriptional targets (**Fig. 1G and 2F**). Unbiased transcriptome analyses via RNA-Seq showed that TGFBR2 expression enriches for mesenchymal signatures and depletes proneural, NPC-like, AC-like and OPC-like signatures in these cell populations (**Fig. 2G**). Consistent with mesenchymal induction, transgenic TGFBR2 induced a shift towards a CD44-high, CD133-low GSC state as determined by western blot and flow cytometry (**Fig. 2H and 2I**). Collectively, these results demonstrate that TGFBR2 is sufficient to induce a mesenchymal shift in GSCs.

### TGFBR2 induces an immunosuppressive ITL signature in mesenchymal GSCs

To identify novel molecular events driven by TGFBR2 signaling in GSCs, we performed an unbiased gene-set enrichment analysis (GSEA) on the genes differentially induced by transgenic TGFBR2. Interestingly, this analysis revealed enrichment of several gene signatures related to the Treg phenotype (**Fig. 3A**). To determine if these gene signature changes were specific to the Treg phenotype, we performed GSEA using gene signatures corresponding to multiple immunosuppressive cell types typically found in the GBM immune TME (i.e., Tregs, myeloid-derived suppressor cells (MDSCs), suppressive M2-like macrophages, and tumor-associated neutrophils) (26–33). The TGFBR2-induced transcriptome was most consistently and significantly associated with the Treg signature, inconsistently associated with the MDSC signature and unassociated with either macrophage or neutrophil signatures (**Fig. 3B**). We identified 362 unique genes upregulated by transgenic TGFBR2 in GBM neurospheres and associated with a Treg state (**Table S1**), consistent with the hypothesis that TGFBR2 mediates an immunosuppressive mesenchymal shift that resembles Treg functionality in GSCs. Tregs reprogram the immune TME by inhibiting anti-tumor immune cell function in a variety of ways including releasing cytokines and proteins and signaling through ligand-receptor interactions and ectoenzymes on the cell surface (34). To identify genes that may play a direct immunosuppressive role in mGSCs, we first queried a scRNA-seq dataset of patient-derived GSCs for genes known to be direct immunosuppressive effectors in Tregs. Using this approach we identified CD274 (PD-L1), NT5E (CD73), ENTPD1 (CD39), LGALS1 (galectin-1), PDCD1LG2 (PD-L2), and TGFB1 as putative immunosuppressive factors in this cell subset **(Fig. S1)**. Of note, we did not detect transcripts for FOXP3, CD25, IL2R, master regulators of Treg development and function, or IL10 and Granzyme B (34). Analysis of clinical GBM transcriptomic datasets (TCGA and Rembrandt) showed a positive correlation between the 6 immunosuppressive genes identified and TGFBR2^High^, CD44^High^ and mesenchymal GBM cell subsets, as defined by Neftel et al and Verhaak et al (25, 35) **(Fig. 3C).** We refer to this gene set as the immunosuppressive Treg-like (ITL) signature.

**Figure 3.**
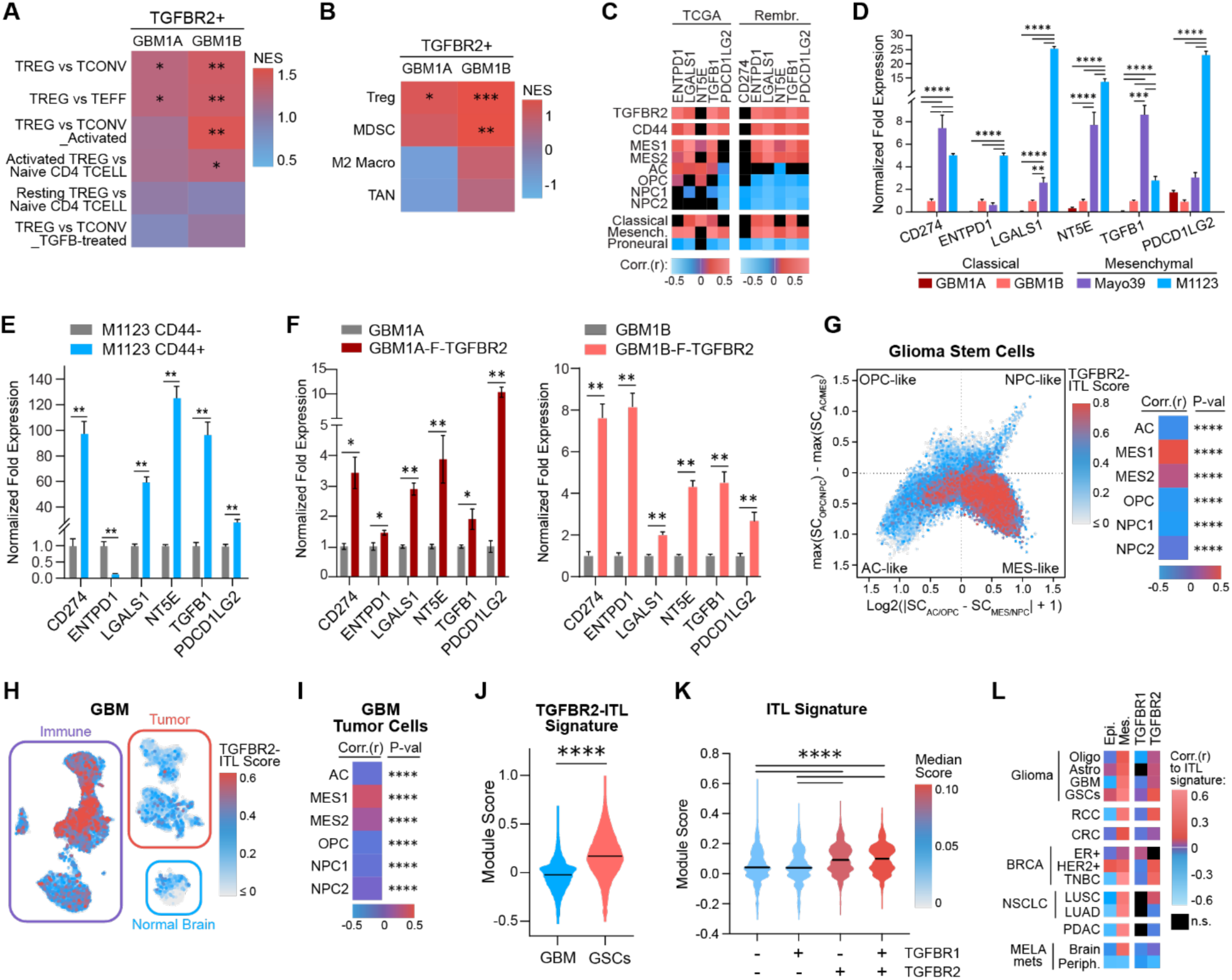
TGFBR2 induces an immunosuppressive Treg-like signature in mesenchymal GSCs. **(A)** Heatmap of GSEA showing enrichment of Treg-related genes in GSCs expressing transgenic TGFBR2. NES = normalized enrichment score. **(B)** Heatmap of GSEA showing enrichment scores for gene signatures for immunosuppressive cell types in GSCs expressing transgenic TGFBR2. MDSC = myeloid-derived suppressor cell; M2 Macro = M2-polarized macrophage; TAN = tumor-associated neutrophil. **(C)** Heatmap showing Pearson’s coefficient values comparing Treg effector genes and TGFBR2, CD44, GBM cell states, and GBM molecular subtypes in clinical GBM specimens. TCGA = TCGA HG-U133A, Rembr. = Rembrandt**. (D)** qRT-PCR analysis measuring expression of Treg effector genes in GBM neurospheres. **(E)** qRT-PCR analysis comparing expression of Treg effector genes in CD44+ versus CD44-GSCs. **(F)** qRT-PCR analysis comparing expression of Treg effector genes in GSCs -/+ transgenic TGFBR2. **(G)** Cell state plot (left) showing expression of the TGFBR2-ITL signature in GBM cells enriched for GSCs derived from 26 patient tumors (36). Heatmap showing Pearson’s coefficient (right) from the same cells showing correlations to the GBM cell states. **(H)** UMAP showing expression of the TGFBR2-ITL signature in scRNA-seq from clinical GBM specimens. Single-cell data from 7 patient GBMs was obtained from Richards et al (36). **(I)** Heatmap showing Pearson’s coefficient from the tumor cells in panel H showing correlations to the GBM cell states. **(J)** Frequency distribution plot of the TGFBR2-ITL gene signature score in tumor cells from clinical GBM specimens compared to patient-derived GSCs. **(K)** Frequency distribution plot of the ITL gene signature score in patient-derived GSCs grouped by -/+ expression of TGFBR1 and TGFBR2. **(L)** Heatmap showing Pearson’s coefficient from a pan-cancer analysis showing correlations between the ITL signature and TGFBR1, TGFBR2 and epithelial (Epi.) & mesenchymal (Mes.) signatures in various cancer types. Oligo = oligodendroglioma, Astro = astrocytoma, RCC = renal cell carcinoma, CRC = colorectal cancer, BRCA = Breast cancer, TNBC = triple-negative breast cancer, NSCLC = non-small cell lung cancer, LUSC = lung squamous cell carcinoma, LUAD = lung adenocarcinoma, PDAC = pancreatic ductal adenocarcinoma, MELA mets = melanoma metastases, Periph. = peripheral, n.s. = non-significant. Data in panels D, E, & F are shown as mean ± SD. Statistical significance was calculated using the nominal p-value in panels A and B, Pearson’s correlation in panels C, G, I & L, one-way ANOVA with Tukey’s post hoc test in panels D, Student’s T-test in panels E & F, Mann-Whitney U-test in panel J, and Kruskal-Wallis test in panel K. *p<0.05, **p<0.01, ***p<0.001, ****p<0.0001.

Consistent with these transcriptomic associations, we measured higher expression of the ITL signature genes in mGSCs compared to classical neurosphere isolates (**Fig. 3D**) and 5 out of 6 genes were enriched in CD44+ cells compared to their CD44-counterparts (**Fig. 3E**). Additionally, we determined that transgenic expression of TGFBR2 is sufficient to induce the mRNA expression of all 6 genes in the ITL signature in 2 distinct classical neurosphere isolates (**Fig. 3F**). As predicted, this TGFBR2-induced ITL (TGFBR2-ITL) gene signature is highly expressed in MES-like patient-derived GBM cells enriched for GSCs (**Fig. 3G**) (36). Critically, this signature was also found to be expressed in neoplastic tumor cells within clinical GBM pathology specimens (**Fig. 3H**) (36) and is specifically enriched in the MES-like GBM cell subsets (**Fig. 3I**). This TGFBR2-driven ITL gene signature has a significantly higher expression in patient GSCs compared to all neoplastic GBM cells within patient tumors (**Fig. 3J**), emphasizing the enrichment of this signature in stem-like cells. Notably, this ITL signature did not correlate with TGFBR1 expression, indicating a distinct role for TGFBR2 in controlling the ITL GSC phenotype (**Fig. 3K**). Furthermore, a pan-cancer bioinformatics analysis revealed a strong association between the 6-gene ITL signature and both mesenchymal signatures and TGFBR2 expression in a variety of systemic cancers (**Fig. 3L**). Together, these results identify a TGFBR2-induced signature of immunosuppressive effector genes, known to be expressed by Tregs, in mesenchymal cancer cells across multiple solid tumor types.

### TGFBR2 inhibition blocks the immunosuppressive GSC phenotype

Our results show that TGFBR2 induces a subset of immunosuppressive effectors associated with Treg function, predicting that TGFBR2 inhibition would reduce the immunosuppressive capacity of GSCs. To test this, we utilized Inducer of TGFBR2 Degradation-1 (ITD-1), a small-molecule TGFBR2 inhibitor that activates proteasome-dependent TGFBR2 protein degradation (37) and GBM cells engineered for doxycycline-induced shRNA-mediated TGFBR2 expression knockdown. Quantitative immunofluorescence analysis confirmed that ITD-1 depleted TGFBR2 protein (**Fig. 4A**) and qRT-PCR showed that shTGFBR2 inhibited TGFBR2 expression in patient-derived mGSCs (**Fig. S2A**). ITD-1 and shTGFBR2 also inhibited expression of the 6-gene ITL signature in patient-derived mGSCs (**Fig. 4B & S2B**). These results show that TGFBR2 signaling is required to maintain expression of the 6-gene ITL signature and predicts that ITD-1 and shTGFR2 will reduce the immunosuppressive phenotype of mGSCs.

**Figure 4.**
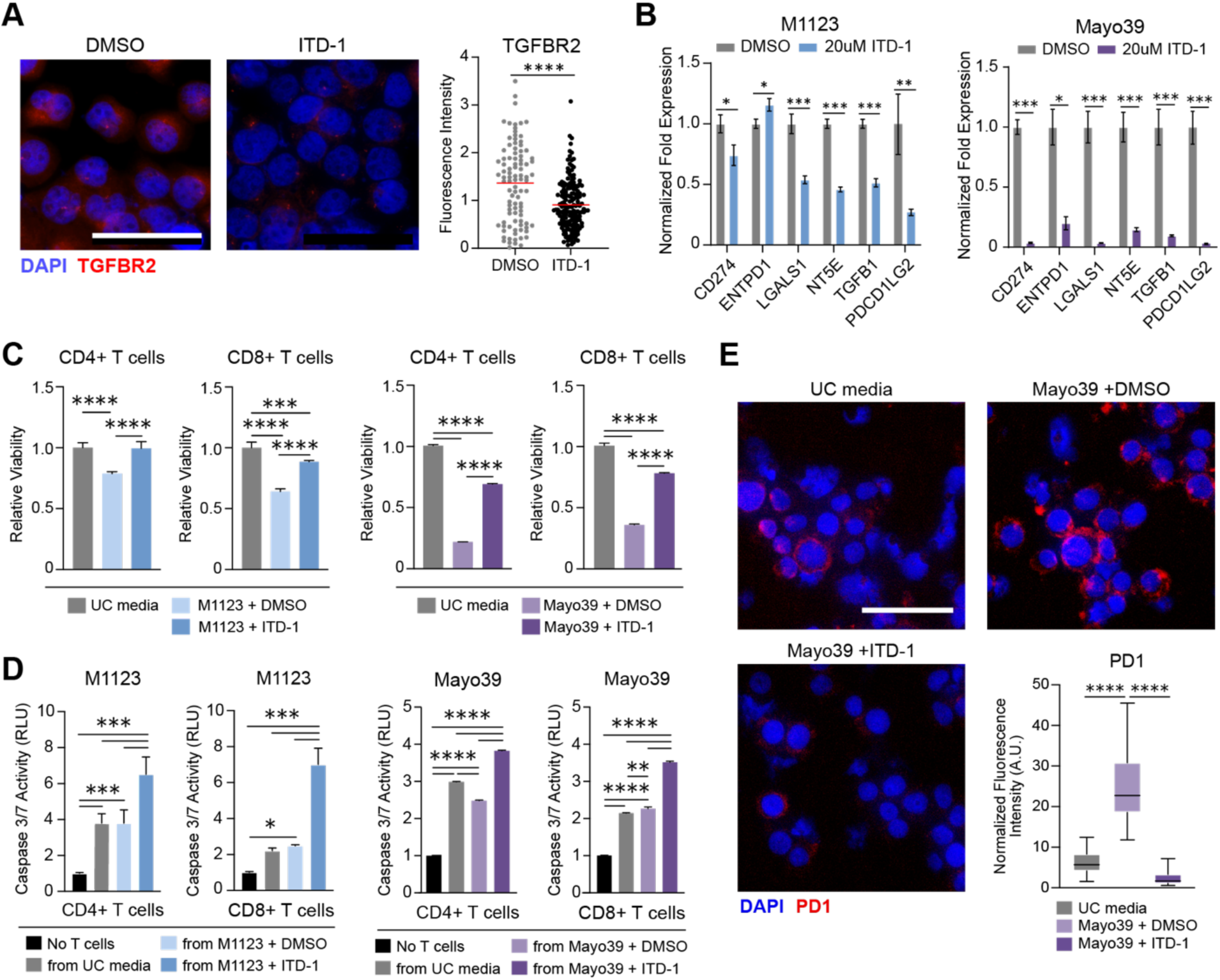
Pharmacological TGFBR2 inhibition attenuates the immunosuppressive phenotype of GSCs. **(A)** Representative immunofluorescence images (left) and quantification (right) showing TGFBR2 expression in M1123 cells 24h after treatment with ITD-1 (20μM) or vehicle control (DMSO). Scale bar = 50μm. **(B)** qRT-PCR analysis showing expression of Treg effector genes 72h following treatment with ITD-1 (20μM) or vehicle control (DMSO). **(C)** CD4+ and CD8+ T cell viability was measured 48h after culture in media conditioned by GSCs treated with ITD-1 or DMSO. CD4+ and CD8+ T cells cultured in unconditioned (UC) media is used as a control. **(D)** Tumor cell death was measured via caspase 3/7 assay 48h after co-culture with CD4+ or CD8+ T cells cultured in media conditioned by GSCs pre-treated with ITD-1 or DMSO as described in Materials & Methods. Tumor cells cultured in the absence of CD4+ or CD8+ T cells is used as a baseline control. **(E)** Representative immunofluorescence images and quantification (lower right) of PD1 expression in CD8+ T cells 48h after culture in media conditioned by GSCs treated with ITD-1 or DMSO. Scale bar = 50μm. PD1 fluorescence intensity was measured using ImageJ’s ‘Analyze Particles’ feature and normalized to total DAPI intensity in multiple fields of view (n=20 per condition). Statistical significance was calculated using Mann-Whitney U-test for panel A, Student’s T-test for panel B, one-way ANOVA with Tukey’s post hoc test for panels C & D and Kruskal-Wallis test for panel E. Data are shown as mean ± SD for all bar graphs. *p<0.05, **p<0.01, ***p<0.001, ****p<0.0001.

We examined the effects of conditioned medium (CM) obtained from mGSCs +/- TGFBR2 inhibition on the viability and tumor cell-killing capacity of PBMC-derived CD4+ and CD8+ T cells. Compared to the effect of unconditioned medium, medium conditioned by DMSO-treated mGSCs significantly reduced the viability of CD4+ and CD8+ T cells. The capacity of CM to inhibit T cell viability was lost if collected from mGSCs pre-treated with ITD-1 or from TGFBR2-knockdown GBM cells (**Fig. 4C & S2C**). CD4+ and CD8+ T cells cultured in CM obtained from mGSCs pre-treated with ITD-1 or from TGFBR2-knockdown GBM cells displayed enhanced tumor cell-killing capacity compared to T cells cultured in either unconditioned medium or in CM from control mGSCs (DMSO-treated or no doxycycline) (**Fig. 4D & S2F**). We further investigated the impact of inhibiting GBM cell TGFBR2 on CD8+ T cell expression of the exhaustion marker PD-1. CM from mGSCs markedly induced CD8+ T cell expression of PD-1, an effect completely abrogated by inhibiting neurosphere cell TGFBR2 with either ITD-1 or shTGFBR2 prior to CM collection (**Fig. 4E& S2D**). Collectively, these results demonstrate that selective targeting of TGFBR2 represses the ITL gene signature in GSCs and inhibits their immunosuppressive effects on CD4+ and CD8+ T cells.

## Discussion

The immunosuppressive TME is a hallmark of GBM and GBMs characterized by a mesenchymal transcriptome contain an especially high proportion of immunosuppressive cells along with having the shortest median survival and reduced sensitivity to standard-of-care therapy relative to other molecular subtypes (38–40). Tumor cells are known modulators of the immune GBM TME with GSCs having potent immunosuppressive capacity (41). Recent studies have highlighted multiple GSC-specific factors responsible for recruiting and polarizing TAMs to a pro-tumor M2-like phenotype (42–44) or suppressing anti-tumor T cell infiltration and function (3, 45). However, the extent to which GSC-intrinsic mechanisms impede anti-tumor immune cell function and the targetable factors responsible are not fully understood. Our lab has recently described how stemness-driving events coordinated by Oct4 and Sox2 enhance tumorigenicity and tumor cell-mediated immunosuppression (5, 21–24). In particular, Oct4/Sox2 upregulate expression of certain immune checkpoint molecules, cytokines, and chemokines in a BRD4-dependent manner, resulting in enhanced T cell apoptosis, Treg infiltration, and immunosuppressive M2-like macrophage polarization (5). We now show that stem cell-reprogramming events initiated by Oct4 and Sox2 induce a mesenchymal transition in GBM cells characterized by activation of TGFBR2 signaling (**Fig. 1 and 2**) which in turn mediates transcriptome changes resembling a Treg state (**Fig. 3 and S1**). Critically, the immunosuppressive phenotype of TGFBR2^High^ mGSCs is blocked by TGFBR2 inhibition, decreasing CD8+ T cell exhaustion and restoring CD4+ and CD8+ T cell tumor cell-killing ability *in vitro* (**Fig. 3, 4, and S2**).

Previous research shows that tumor cells are capable of co-opting developmental pathways associated with non-neoplastic cells to support tumor growth and therapeutic resistance (46). In GBM, tumor cells have demonstrated mimicry of vascular cells (47) and neuronal- and glial-progenitor cells (48, 49). Moreover, GSCs can acquire a myeloid-like transcriptional profile following repeated immune exposure, facilitating immune evasion and promoting infiltration and polarization of pro-tumor myeloid cells (50). We now show that TGFBR2 associates with and activates a Treg-like state as GSCs become more mesenchymal and immunosuppressive **(Fig. 2, 3, 4, and S1).** This recapitulates to some extent the capacity of TGFβ signaling to differentiate naïve T cells into an induced Treg state (51) and further delineates its role in acquisition of immunosuppressive cell states. Similar to previous reports describing Foxp3-CD103+ T cell-derived induced Tregs (52), the absent expression of canonical Treg markers Foxp3 and CD25 in the GSC-derived induced Treg-Like cells described here suggests that GSCs do not transdifferentiate into Tregs in response to TGFBR2 but instead co-opt certain mechanisms utilized by Tregs to exert immunosuppressive behavior. Among the multitude of genes induced by TGFBR2, we identified a 6-gene signature comprised of putative direct effectors of the immunosuppressive GSC phenotype (**Fig. 4C-H**). These include NT5E (CD73) and ENTPD1 (CD39) which are involved in immunosuppressive adenosine signaling (53), LGALS1 (galectin-1) and TGFB1 which encode for anti-inflammatory cytokines (54, 55), and CD274 (PD-L1) and PDCD1LG2 (PD-L2) which bind to PD-1 to initiate an inhibitory signaling pathway in anti-tumor immune cells (56). Notably, this molecular phenotype correlates with previously defined cell states in GBM associated with immune cell interaction and an immunosuppressive TME (**Fig. 3**) (25, 57). We also show that the correlation of this immunosuppressive signature with TGFBR2 and the mesenchymal state is conserved in a variety of systemic cancers (**Fig. 3L**), suggesting that findings from this study may be applicable to multiple tumor types.

Although blocking TGFβ signaling is widely viewed as a promising anti-tumor strategy, efforts in GBM have mainly focused on blocking TGFBR1 activity without successful clinical translation (58–60). This might be explained by our current results specifically identifying TGFBR2 as the driver of GSC-derived induced Treg-like cells and associating their immunosuppressive signature with TGFBR2 but not TGFBR1 expression in clinical specimens across multiple cancers, uncovering previously unrecognized TGFBR2 dependencies in cancer **(Fig. 3K and 3L)**. Currently, the only TGFBR2-specific drug that has been tested in a clinical setting is IMC-TR1 (LY3022859), an anti-TGFBR2 monoclonal antibody, which showed no efficacy in Phase 1 trial for advanced solid tumors (61). Despite the limited efficacy of these inhibitors, alternative approaches employing combinations with immunotherapies still hold promise (62). We show that blocking TGFBR2 via an shRNA or a selective inhibitor, ITD1 (37), reduces the expression of this gene signature in mGSCs (**Fig. 4C and S2B**) and we demonstrate through T cell culture assays that mGSCs expressing this ITL transcriptional signature reduce CD4^+^ and CD8^+^ T cell viability and function, which can be rescued by blocking TGFBR2 signaling (**Fig. 4 and S2**).

In summary, we describe a mechanism by which stem cell-driving events coordinate the transition to a mesenchymal-like GSC state through activation of TGFBR2 signaling (**Fig. 5A**). In turn, TGFBR2 induces an immunosuppressive GSC phenotype reminiscent of Tregs that allows GSCs to inhibit T cell function by decreasing proliferation capacity and inducing exhaustion (**Fig. 5B**). Blocking TGFBR2 signaling in mGSCs, through both molecular and pharmacological methods, countered this GSC-mediated immunosuppression, predicting that TGFBR2 blockade could cooperate with current immunotherapy to enhance anti-tumor effects in GBM.

**Figure 5.**
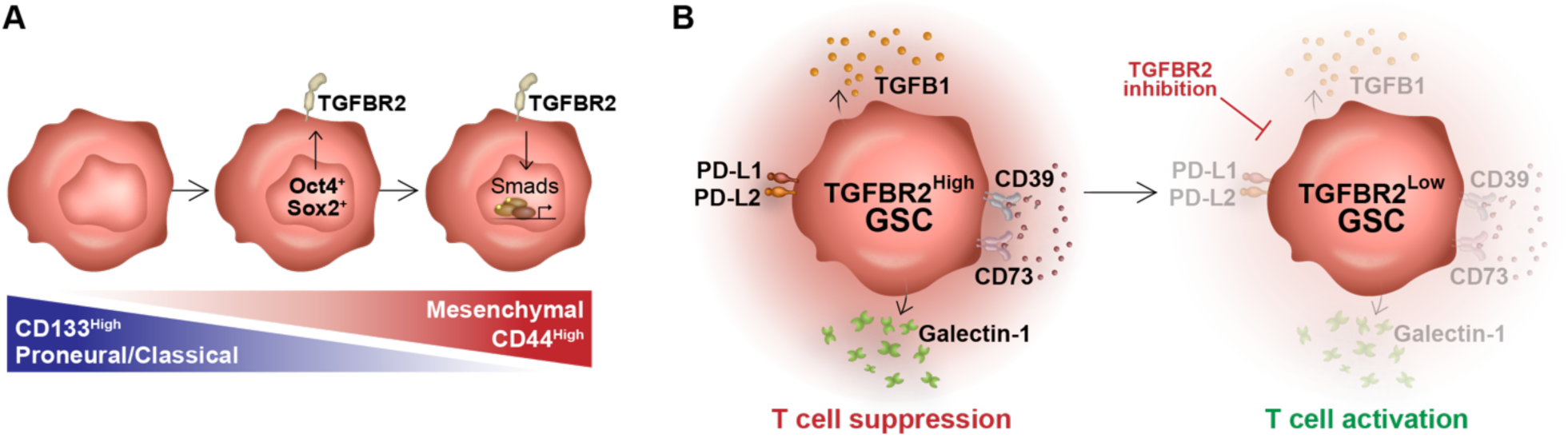
Graphic Summary. **(A)** Glioma stem cells in a proneural, CD133^high^ state can transition to a mesenchymal, CD44^high^ state following co-expression of the reprogramming transcription factors Oct4 and Sox2. Oct4 and Sox2 induce expression of TGFBR2, activating downstream signaling and promoting a mesenchymal state. **(B)** High TGFBR2 expression leads to upregulated expression of immunosuppressive effector genes to generate a T cell-suppressing GSC phenotype. TGFBR2 inhibition attenuates expression of these effectors, allowing for T cell activation.

## Materials & Methods

### Human cell culture

All GSCs used in this study were derived from newly diagnosed GBMs (IDH-wildtype) and cultured in serum-free conditions using Stemline Neural Stem Cell Expansion Medium (Sigma-Aldrich, St. Louis, MO, USA) supplemented with 20ng/mL epidermal growth factor and 10ng/mL fibroblast growth factor. The classical-like patient-derived neurosphere lines, GBM1A and GBM1B, were originally derived and characterized by Vescovi and colleagues (63). The mesenchymal-like patient-derived GBM xenograft cell line, Mayo39, was obtained from the Mayo Clinic (Rochester, MN) (64) and enriched for stem-like cells by culturing in Stemline Neural Stem Cell media prior to use in experiments. Low-passage patient-derived mesenchymal GSCs, M1123, were a kind gift from Dr. Nakano at The Ohio State University (65). HEK293FT cells were obtained from the ATCC and grown in Dulbecco’s modified Eagle medium (DMEM) supplemented with 10% FBS (Fetal Bovine Serum, Thermo Fisher, Waltham, MA, USA). Prior to experimentation, all cells were tested for mycoplasma contamination and authenticated via STR profiling.

### Lentivirus generation and cell transduction

The transgenic cell lines used in this study were generated with the second-generation lentiviral system according to Addgene protocols, using the psPAX2 packaging plasmid and pMD2.G envelope plasmid (Addgene, Watertown, MA, USA). The lentiviral packaging/envelope plasmids and the transgene vector (**Supplementary Table 2**) were co-transfected into HEK293FT cells using the Lipofectamine 3000 kit (Thermo Fisher) according to the manufacturer’s recommendations. The next morning, sodium butyrate (Cayman Chemical, Ann Arbor, MI, USA) was added to transfected cells at a final concentration of 10mM to enhance the viral titer. After 48h, lentiviral particles in the supernatant were concentrated using Lenti-X concentrator solution (Takara Bio, San Jose, CA, USA) and resuspended in 1mL PBS to transduce cells. GSCs were infected overnight with the lentivirus particles plus 1ug/mL polybrene. The next morning, cells were replated in fresh neurosphere media.

### Western blot analysis

To measure protein expression, cells were lysed in RIPA buffer (Sigma-Aldrich) plus protease inhibitors (Sigma-Aldrich #P8340) and phosphatase inhibitors (Sigma-Aldrich #P5726) for 30min on ice. Protein was purified by centrifugation and quantified by Bradford protein assay. Equal quantities of protein were loaded per sample (40-80ug) and resolved on a NOVEX 4-12% or 4-20% Tris-glycine gradient gel (Thermo Fisher) using the Thermo Fisher Mini Gel Tank system. Protein was then transferred onto an Amersham Protran nitrocellulose membrane (GE Healthcare, Chicago, IL, USA) using the Bio-Rad Mini Protean 3 Cell system. The membrane was blocked for 1h in Li-COR Intercept blocking buffer before primary antibodies (**Supplementary Table 3**) were added and incubated overnight at 4°C. Membranes were then washed and incubated with infrared-labeled secondary antibodies (Li-COR Biosciences, Lincoln, NE, USA) prior to quantification using the Odyssey CLx Infrared Imager (Li-COR Biosciences). Densitometry analysis was performed using Image Studio software from Li-COR imaging systems. Protein expression was normalized to the loading control (e.g., GAPDH).

### qRT-PCR analysis

To measure gene expression, total RNA was extracted from cells using the RNeasy Mini Kit (Qiagen, Germantown, MD, USA) and converted to cDNA by reverse-transcribing 500ng-1ug of RNA using MuLV Reverse Transcriptase and Oligo (dT) primers (Applied Biosystems, Waltham, MA, USA). Expression was measured using the Power SYBR Green PCR kit (Applied Biosystems) and quantified using a Bio-Rad CFX96 Real-Time Detection System and accompanying software. Samples were run in triplicates and signal was normalized to 18S RNA. Primer sequences are provided in **Supplementary Table 4**.

### Cell viability and cell death assays

To quantify cell viability and cell death, cells were split evenly and incubated at room temperature (RT) in either Cell Titer Glo (Promega, Madison, WI, USA) or Caspase 3/7 Glo (Promega) at a 1:1 (vol/vol) ratio to cell media. After 30min, luminescence was measured using the SpectraMax M5 Multimode Plate Reader (Molecular Devices, San Jose, CA, USA) and quantified using SoftMax Pro 7 software. Cell viability was calculated as the ratio of Cell Titer Glo to Caspase 3/7 Glo signal, whereas cell death is represented by the inverse value.

### Flow cytometry and FACS

To quantify cell proportions, GSCs were dissociated into single cells and incubated with the appropriate fluorescently labeled antibodies (**Supplementary Table 3**) following the manufacturer’s recommendations (Miltenyi Biotec, Gaithersburg, MD, USA). Quantification was performed using the Muse® Cell Analyzer (Sigma-Aldrich) and gated for cell size and fluorescence signal. To sort GSCs into CD44-high and -low fractions, GSCs were processed the same as above and sorted using the Beckman Coulter’s MoFlo Astrios EQ cell sorter.

### RNA sequencing

RNA-Seq libraries were constructed from messenger RNA (mRNA) purified from total RNA using poly-T oligo-attached magnetic beads. After fragmentation, the first strand cDNA was synthesized using random hexamer primers, followed by the second strand cDNA synthesis using dUTP. The library was checked with Qubit and real-time PCR for quantification and bioanalyzer for size distribution detection. Quantified libraries were pooled and sequenced on Illumina platforms followed by clustering of the index-coded samples according to the manufacturer’s instructions. After cluster generation, the library preparations were sequenced on an Illumina platform and paired-end reads were generated. Index of the reference genome (i.e. hg38) was built and reads were aligned to the reference genome using Hisat2 v2.0.5. Differential expression analysis of two conditions/groups (two biological replicates per condition) was performed using the DESeq2 R package (1.20.0).

ScRNA-seq from GBM1A and GBM1A-Oct4/Sox2+ cells was performed using the 10X Genomics Chromium v2 platform according to standard protocol. Reads were sequenced using the Illumina NovaSeq system and aligned using the hg38 genome. Count matrices were generated using CellRanger. Data processing was conducted using the Seurat v5 package in R Studio (66). Low quality cells, defined as number of features and/or counts < 500 and percentage of mitochondrial reads > 20%, were excluded and counts were normalized using the *SCTransform()* function prior to downstream analyses. Batch correction was performed using Harmony (67). Dimensionality reduction was performed using the first 20 principal components when applicable.

Expression scores for gene signatures in bulk RNA-seq samples were calculated using single-sample gene-set enrichment analysis (**Supplementary Table 5**) (68). In scRNA-seq, gene signatures for 6 GBM cellular states (MES1, MES2, AC, OPC, NPC1, and NPC2) were obtained from Neftel et al. (25) and scores were calculated using the *AddModuleScore()* function in Seurat with the following parameters: nbins = 30 and ctrl = 100. The MES and NPC module scores were calculated by averaging the MES1/MES2 and NPC1/NPC2 values, respectively. To generate the cell state plot, the y-axis coordinate for each cell was calculated as y = max(SCopc, SCnpc) - max(SCac, SCmes) where SC = state module score for a given cell. A positive y-axis value indicates OPC/NPC lineage while a negative value indicates AC/MES lineage. The x-axis coordinate was then calculated using the following formulas, depending on lineage. For OPC/NPC lineage, x = log2(ABS(SCopc - SCnpc) +1). For AC/MES lineage, x = log2(ABS(SCac - SCmes) +1). Custom R scripts used to generate data figures are available upon request.

### Analysis of publicly available GBM expression data

TCGA (HG-U133A) and Rembrandt clinical and transcriptional data from patient glioma and non-tumor brain specimens were obtained from the GlioVis data portal (http://gliovis.bioinfo.cnio.es/). RNA-seq data from patient-derived xenograft (PDX) cell lines and patient-derived GSC lines were obtained from cBioPortal and GSE119776, respectively. Patient GBM and GSC scRNA-seq data was obtained from the Broad Institute Single Cell Portal (www.singlecell.broadinstitute.org) under study SCP503. A list of the scRNA-seq datasets used for the pan-cancer analysis shown in **Figure 3L** can be found in the **Supplementary Table 6**. Count matrices for scRNA-Seq analysis were processed as described above and cell annotations, normalized gene expression and UMAP coordinates from the original publications were used for all other downstream analyses.

### Isolation and activation of PBMC-derived T cells and Immune cell co-culture assay

CD4+ and CD8+ T-cells were isolated from patient-derived PBMCs (69) using MOJO anti-CD4 and anti-CD8 bead isolation kits (BioLegend, San Diego, CA, USA), respectively. T cells were then activated via anti-CD28/CD3 Dynabeads™ (Gibco, Thermo Fisher) and recombinant IL2 (Peprotech, Thermo Fisher; 25U per 1×10^5 cells). Activated cells were cultured in RPMI-1640 media containing 2mM L-glutamine, 10mM HEPES, 1mM sodium pyruvate, 4500mg/L glucose, and 1500mg/L sodium bicarbonate, and supplemented with 10% fetal bovine serum, as recommended by the ATCC.

The effect of GSCs on CD4+ and CD8+ T-cells was assessed by culturing the T cells in GSC-conditioned media (CM) for 48-72h. To generate the CM, GSCs were cultured in neurosphere medium +/- ITD-1 (20 μM) or in GSC medium +/- doxycycline (1 μg/mL) to induce shTGFBR2 for 48h and 5 days, respectively. GSCs were then rinsed and replated at equal densities (∼5×10^5^ cells/mL) in fresh neurosphere medium (lacking ITD-1 or doxycycline) for 48h. CM was then collected and added to immune cells plated at 100-200K cells/well in 24-well cell culture plates at a 1:1 ratio (vol/vol) to immune cell media. Cells were collected to quantify cell viability and measure gene expression changes. The cytotoxic effects of T-cells on GSCs were assessed by plating GSCs on laminin-coated plates at equal densities (∼100-200K cells/well) and co-culturing with CD4^+^ or CD8^+^ T cells in T cell media for 48h. GSC death was analyzed after removing T cells and washing wells with phosphate-buffered saline (PBS).

### Immunofluorescence imaging

To quantify TGFBR2 knockdown (**Fig. 4A**) and T cell exhaustion (**Fig. 4E and S2E**), cells were collected, counted, and then spun onto microscope slides at a density of 100K cells per spot using cytospin technology. Cells were fixed for 20min with a 1% paraformaldehyde solution and then washed with PBS before blocking for 1h with PBS containing 1% bovine serum albumin (BSA; Sigma-Aldrich). Fluorescence-conjugated primary antibodies (**Supplementary Table 3**) were added onto cells (1:200-1:500 in 1% BSA-PBS) and incubated at 4°C overnight. The next day, cells were washed with PBS and coverslips were mounted with Prolong Gold Antifade plus DAPI mounting media (Cell Signaling, Danvers, MA, USA). Slides were imaged with Leica DMi8 Thunder Imager Live Cell microscope and fluorescence signal was quantified using ImageJ software with background noise removal (NIH). Protein expression was calculated relative to DAPI signal in each field-of-view (40X magnification).

### Statistical analyses

All experiments were performed in triplicates and repeated at least twice in each cell model (N ≥ 6). PRISM GraphPad 10 was used to perform all the statistical analyses presented. Two group comparisons were analyzed for variation and significance using a two-tailed, type 1 t-test and p-values lower than 0.05 were considered significant and symbolized by an asterisk in the graphs. One-way ANOVA and Tukey post hoc tests were used to analyze the relationships when comparing multiple experimental groups with p-values lower than 0.05 considered to be statistically significant. All data shown are representative of mean ± SD of triplicate results unless otherwise specified.

## Acknowledgements

The authors would like to thank the following organizations for financial support: Johns Hopkins University Provost’s Undergraduate Research Award (HK); Clinical Translation Core at the Kennedy Krieger Institute (5P50HD103538-02); and the United States NIH grants R01NS096754 (JL), R01NS120949 (HLB), and F99CA284254 (ALJ)

## Notes

### Competing Interest Statement

The authors have declared no competing interest.

