## Supplemental Figures and Tables for "TGFBR2^High^ mesenchymal glioma stem cells phenocopy regulatory T cells to suppress CD4+ and CD8+ T cell function"

### Supplemental Data

#### Supplemental Figures

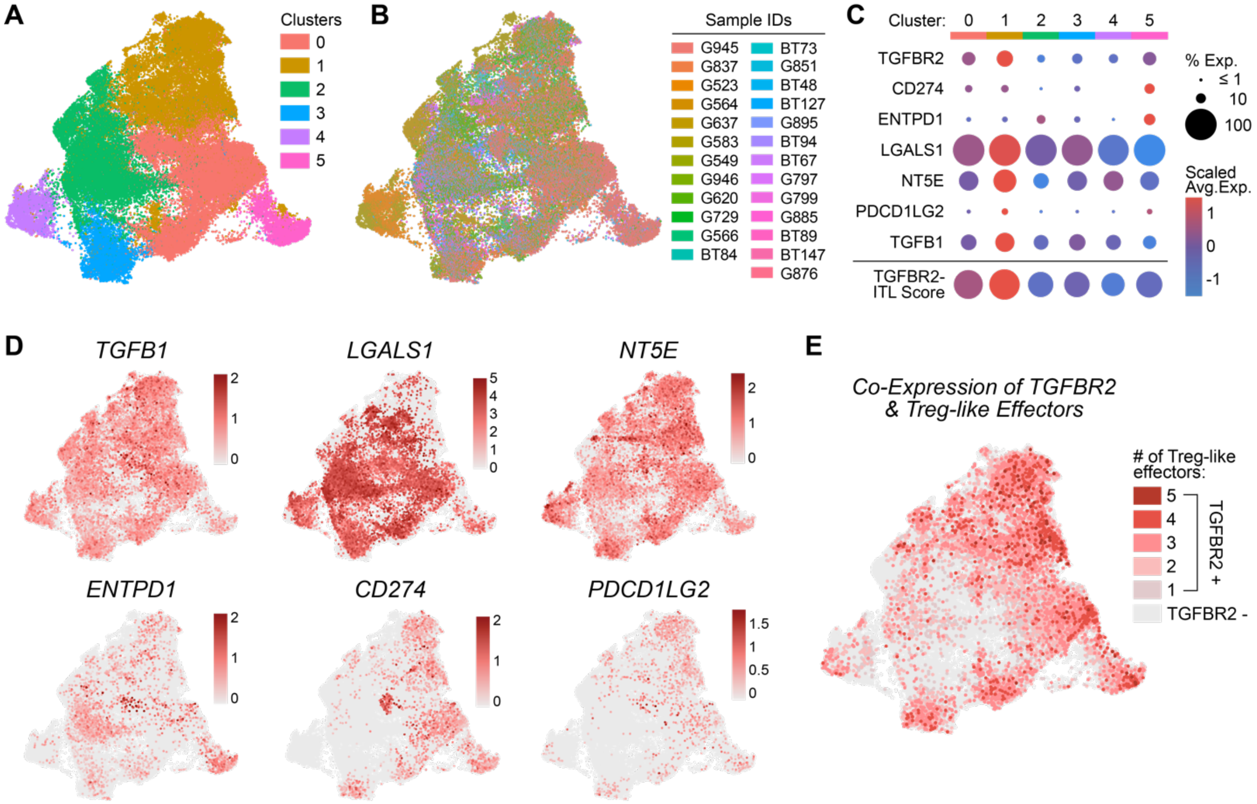


**Supplemental Figure 1. Canonical regulatory T cell effector genes are expressed in TGFBR2+ patient-derived GSCs. (A)** UMAP showing clustering of patient-derived GSCs. **(B)** UMAP showing distribution of patient samples following Harmony batch correction. **(C)** Dot plot showing expression levels of TGFBR2, regulatory T cell genes, and the TGFBR2-ITL gene signature in cell clusters from patient-derived GSC scRNA-seq. **(D)** UMAPs showing expression of individual Treg-related effector genes in patient-derived GSCs. **(E)** UMAP showing degree of co-expression of Treg-related effectors with TGFBR2 in patient-derived GSCs.

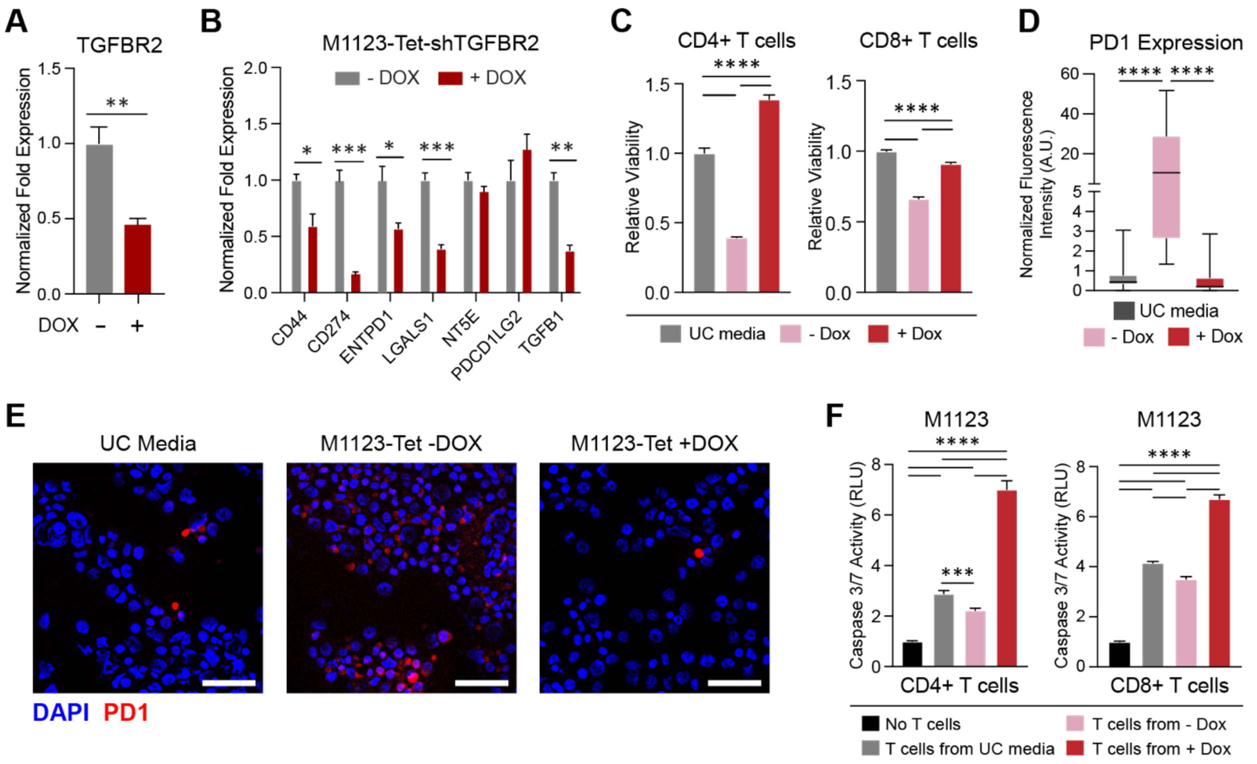


**Supplemental Figure 2. ShRNA-mediated TGFBR2 knockdown in GSCs attenuates the immunosuppressive ITL phenotype. (A)** qRT-PCR analysis showing knockdown of TGFBR2 expression in M1123-Tet-shTGFBR2 cells 72h following doxycycline (Dox) treatment. **(B)** qRT-PCR analysis showing knockdown of Treg effector genes in M1123-Tet-shTGFBR2 cells 5-days following Dox. **(C)** T cell viability assays showing an increase in cell viability when CD4+ or CD8+ T cells are cultured in media conditioned by M1123 cells with TGFBR2 inhibition (+ Dox) compared to control (-Dox). UC media = unconditioned media. **(D)** Quantification and **(E)** representative immuno-fluorescence images of PD1 expression in CD8+ T cells following culture in media conditioned by M1123-Tet-shTGFBR2 cells -/+ Dox. Quantification was performed from immunofluorescence images and calculated in multiple fields of view (40X magnification, n=20-30 per condition) and normalized to total DAPI signal. Scale bar = 50µm. **(F)** Tumor cell death assays showing enhanced T cell-mediated GSC death following culture in media conditioned by GSCs with TGFBR2 inhibition (+ Dox) compared to controls. Statistical significance was calculated using Student's T-test for panels A & B, one-way ANOVA with Tukey's post hoc test for panels C & F, and Kruskal-Wallis test for panel D. Data are shown as mean ± SD for all bar graphs. *p<0.05, **p<0.01, ***p<0.001, ****p<0.0001.

#### Supplemental Tables

| | | | | | | | |
|---|---|---|---|---|---|---|---|
| ABCC1 | CEP55 | FRMD4B | LAMC1 | POFUT1 | SCAMP1 | SOX3 | TNFSF10 |
| ADAMTS6 | CERK | FRRS1L | LAPTM4A | POLE | SDC4 | SPAG17 | TNFSF12 |
| AGPAT4 | CHORDC1 | FTX | LINC00310 | PPM1B | SEC24A | SPATA22 | TOP3A |
| ALCAM | CHTF18 | FUCA2 | LNPK | PPP1R21 | SEC24D | SPCS2 | TOR1AIP1 |
| ANGPTL1 | CLDN12 | FYN | LRIG1 | PPP1R26-AS1 | SELENOM | SPDYA | TP53 |
| ANKRD13C | CLDND1 | GALM | LRRC8D | PPP6R3 | SEMA5A | SPIN4 | TP53I13 |
| ANKS1B | CLIC1 | GATA1 | LRRCC1 | PRDM1 | SEMA6A | SSR2 | TPH1 |
| ANXA2 | COL1A2 | GBP2 | LYN | PRDM5 | SEMA7A | STC2 | TPR |
| ANXA2P2 | COL7A1 | GBP4 | MAGI1 | PRELID3B | SERPINC1 | STIM2 | TPRG1 |
| ANXA4 | CPD | GCA | MATN2 | PRKRA | SETD6 | STK32A | TRIM16 |
| ANXA5 | CPOX | GCNT1 | MBNL3 | PRNP | SF3B4 | STON1 | TRMO |
| APIP | CRTAP | GGH | MEGF9 | PROS1 | SGSH | SUCO | TRPC3 |
| ARHGAP20 | CSGALNACT2 | GLB1 | MEIS2 | PSEN1 | SH2D1A | SURF4 | TSPAN6 |
| ARL6IP1 | CSRP2 | GLCE | MGME1 | PSRC1 | SHTN1 | SVIP | TTC12 |
| ARMCX4 | CTNS | GLIPR1L1 | MIR22HG | PTBP2 | SIPA1L2 | SWAP70 | TTC39C |
| ARMCX5 | CXCL3 | GNB5 | MPRIP | PTPN7 | SKI | TANK | TXNDC16 |
| ATAD2 | CYB561 | GOLGA8B | MREG | PTPRJ | SKIL | TBK1 | UBE2J1 |
| ATG10 | CYBRD1 | GOSR1 | MT1E | RAB11FIP2 | SLC12A2 | TBRG1 | UBE4B |
| ATL3 | DDX5 | GPR19 | MTERF1 | RAB2B | SLC15A3 | TCEAL8 | UGGT1 |
| ATP11A | DMD | GPR62 | MYL9 | RAD51B | SLC16A1 | TCN2 | USP11 |
| ATP1B1 | DNAH7 | GPR85 | MYO3B | RALB | SLC18B1 | TDRD7 | USP48 |
| ATP2B4 | DNAJC1 | GSAP | NABP1 | RANBP9 | SLC22A1 | TEAD1 | UTRN |
| B2M | DNAJC12 | GUCY1B1 | NDFIP2 | RANGRF | SLC25A24 | TGFBR1 | UTS2B |
| B4GALT4 | DOK2 | HDAC4 | NFKBIZ | RBL1 | SLC25A46 | TGM2 | VAMP3 |
| B9D1 | DYNLL1 | HERPUD1 | NR4A2 | RBPMS | SLC27A2 | THRB | VASN |
| BARD1 | EBI3 | HM13 | NT5E | RDX | SLC35D1 | TIFA | VEZT |
| BECN1 | ECM1 | HSPA1A | NUP155 | REEP3 | SLC39A7 | TINF2 | VPS54 |
| BRCA1 | EGR2 | IBTK | OPTN | REEP5 | SLC44A1 | TIRAP | WDR33 |
| BUB1B | ELL2 | ICAM1 | PCDH11X | RENBP | SLC44A3 | TLCD1 | WFS1 |
| C1QTNF12 | ELOVL5 | IDS | PDCD1LG2 | RHBDL2 | SLC45A1 | TLL2 | WLS |
| C3orf80 | ENTPD1 | IER5 | PDE8A | RHCG | SLC4A11 | TLR4 | WWP1 |
| C6orf136 | EPS15 | IFT80 | PHACTR2 | RNF113A | SLC4A9 | TLR7 | YIPF2 |
| CAPN5 | ERBIN | IGF2R | PHIP | RNF167 | SLC5A3 | TMED3 | YWHAE |
| CASP1 | ERI1 | IL10RB | PHKB | RNF19A | SLC8B1 | TMEM109 | ZC2HC1A |
| CASP3 | ESS2 | IL18 | PHLDB2 | ROBO1 | SLC9A8 | TMEM158 | ZCCHC18 |
| CASP8 | F2RL2 | IL2RB | PHTF2 | ROS1 | SMCHD1 | TMEM208 | ZCCHC8 |
| CCDC50 | FAAP24 | IL7 | PIAS3 | RPE65 | SNAI2 | TMEM65 | ZFAND5 |
| CD200 | FAM174A | IQGAP1 | PIGX | RPL39L | SNHG12 | TMEM70 | ZFAND6 |
| CD274 | FAM174B | IQGAP3 | PIK3AP1 | RRM1 | SNORA65 | TMLHE | ZFYVE16 |
| CD38 | FAR1 | ITGA6 | PIP5K1B | RUBCNL | SNRNP40 | TMTC3 | ZIC1 |
| CD58 | FCHO2 | ITGAE | PIWIL4 | RUNX2 | SNX16 | TNF | ZNF131 |
| CD81 | FES | JADE1 | PLAGL1 | S100A16 | SNX3 | TNFAIP3 | ZNF430 |
| CD83 | FGL2 | KCNMB4 | PLP2 | S100A4 | SOCS2 | TNFRSF18 | ZNF432 |
| CDHR3 | FHL2 | KIF5C | PLS3 | SAMD8 | SOD2 | TNFRSF1B | ZNF644 |
| CDKN2B | FMNL2 | KLHL2 | PLXND1 | SAR1B | SOX2-OT | TNFRSF9 | ZSCAN26 |
| CEP162 | FNBP1L | | | | | | |

**Supplementary Table 1.** Treg-related genes induced by transgenic TGFBR2 in GSCs.

| **Lentiviral Vectors** | | |
|---|---|---|
| *Construct* | *Manufacturer* | *Catalog no.* |
| pEZX-Flag-TGFBR2 | GeneCopoeia | EX-Z4152-Lv242 |
| pTRIPz-shTGFBR2 | Dharmacon | V3THS_406963 |
| pLM-vexGFP-Oct4 | Addgene | #22240 |
| pLM-mCitrine-Sox2 | Addgene | #23242 |

**Supplementary Table 2.** Lentiviral constructs used in this study for stable cell line generation.

| **Antibodies** | | | |
|---|---|---|---|
| *Target* | *Application* | *Manufacturer* | *Catalog no.* |
| GAPDH | Western Blot | Santa Cruz Biotechnology | Sc-47724 |
| Slug | Western Blot | Cell Signaling Technology | 9585T |
| Vimentin | Western Blot | Cell Signaling Technology | 5741T |
| CD44 | Western Blot | Cell Signaling Technology | 156-C311 |
| CD133 | Western Blot | GeneTex | GTX102109 |
| TGFBR2 | Western Blot | GeneTex | GTX129909 |
| Phospho-TGFBR1 | Western Blot | Invitrogen | PA5-40298 |
| Flag | Western Blot | Sigma-Aldrich | F1804 |
| CD44 | Flow Cytometry | Miltenyi Biotec | REA690 |
| CD133 | Flow Cytometry | Miltenyi Biotec | 293C3 |
| TGFBR2 | Immunofluorescence | R&D Systems | FAB241A |
| CD279 (PD1) | Immunofluorescence | Miltenyi Biotec | REA1165 |

**Supplementary Table 3.** List of antibodies used in this study.

| **qRT-PCR Primers** | | |
|---|---|---|
| *Target Gene* | *Forward* | *Reverse* |
| 18S | ACAGGATTGACAGATTGATAGCTC | CAAATCGCTCCACCAACTAAGAA |
| TGFBR2 | GTAGCTCTGATGAGTGCAATGAC | CAGATATGGCAACTCCCAGTG |
| CD274 (PDL1) | TGGCATTTGCTGAACGCATTT | TGCAGCCAGGTCTAATTGTTTT |
| ENTPD1 | AGGTGCCTATGGCTGGATTAC | CCAAAGCTCCAAAGGTTTCCT |
| LGALS1 | TCGCCAGCAACCTGAATCTC | GCACGAAGCTCTTAGCGTCA |
| NT5E | GCCTGGGAGCTTACGATTTTG | TAGTGCCCTGGTACTGGTCG |
| PDCD1LG2 (PDL2) | CTCGTTCCACATACCTCAAGTCC | CTGGAACCTTTAGGATGTGAGTG |
| TGFB1 | TCCTGGCGATACCTCAGCAA | GCCGGTAGTGAACCCGTTGAT |
| CD44 | CTGCCGCTTTGCAGGTGTA | CATTGTGGGCAAGGTGCTATT |
| ANXA4 | GGAGGTACTGTCAAAGCTGCT | GGCAAGGACGCTAATAATGGC |
| IL18 | CAGATCGCTTCCTCTCGCAA | CCAGGTTTTCATCATCTTCAGCTAT |
| SNAI2 | TGCGATGCCCAGTCTAGAAA | AGAAAAAGGCTTCTCCCCCGT |
| F11R | GTGCCTACTCGGGCTTTTCTT | GTCACCCGGTCCTCATAGGAA |
| PDLIM1 | CCCAGCAGATAGACCTCCAG | TCTGAGCTTCCAAGTGTGTCATA |

**Supplementary Table 4.** Primer sequences used in this study for qRT-PCR analyses.

| **Gene set accessions** | | |
|---|---|---|
| *Gene set* | *No. of Genes* | *Source* |
| Smad2/3 Transcriptional Targets | 843 | GSEA-MSigDB; M2356 |
| Verhaak_Mesenchymal | 216 | GSEA-MSigDB; M2122 |
| Verhaak_Classical | 161 | GSEA-MSigDB; M2121 |
| Verhaak_Proneural | 177 | GSEA-MSigDB; M2115 |
| TREG vs TCONV | 199 | GSEA-MSigDB; M4650 |
| TREG vs TEFF | 200 | GSEA-MSigDB; M3673 |

| | | |
|---|---|---|
| TREG vs TCONV_Activated | 200 | GSEA-MSigDB; M5690 |
| Activated TREG vs Naïve CD4 TCELL | 198 | GSEA-MSigDB; M3550 |
| Resting TREG vs Naïve CD4 TCELL | 198 | GSEA-MSigDB; M3542 |
| TREG vs TCONV_TGFB-treated | 199 | GSEA-MSigDB; M5692 |
| Treg (Regulatory T cell) | 321 | doi: 10.1016/j.humimm.2015.12.004 |
| MDSC (Myeloid-Derived Suppressor Cell) | 219 | doi: 10.1126/sciimmunol.aay6017 |
| M2 Macro (M2 Macrophage) | 159 | doi: 10.1038/s41598-020-73624-w |
| TAN (Tumor-Associated Neutrophil) | 200 | doi: 10.1038/s41590-022-01311-1 |
| Pan-cancer Epithelial | 25 | doi: 10.1158/1078-0432.CCR-15-0876 |
| Pan-cancer Mesenchymal | 52 | doi: 10.1158/1078-0432.CCR-15-0876 |

**Supplemental Table 5.** Source information for gene sets used in this study.

| **Publicly Available scRNA-seq Datasets** | | |
|---|---|---|
| *Tumor Type* | *Depository & Accession* | *Publication* |
| Glioma | GEO; GSE182109 | doi: 10.1038/s41467-022-28372-y |
| Renal Cell Carcinoma | TISCH2; ID: T020120; KIRC_GSE171306 | doi: 10.1073/pnas.2103240118 |
| Colorectal Cancer | TISCH2; ID: T020099; CRC_GSE166555 | doi: 10.15252/emmm.202114123 |
| Breast Cancer | TISCH2; ID: T010014; BRCA_GSE176078 | doi: 10.1038/s41588-021-00911-1 |
| Non-Small Cell Lung Cancer | TISCH2; ID: T020140; NSCLC_GSE148071 | doi: 10.1038/s41467-021-22801-0 |
| Pancreatic Ductal Adenocarcinoma | TISCH2; ID: T010066; PAAD_CRA001160 | doi: 10.1038/s41422-019-0195-y |
| Melanoma Metastases | GEO; GSE185386 | doi: 10.1016/j.cell.2022.06.007 |

**Supplemental Table 6.** Source information for publicly available single-cell RNA-sequencing datasets used in this study.
